# Chemical composition of essential oil from *Folium nelumbinis* and its antioxidant activity

**DOI:** 10.1101/419945

**Authors:** Xiaoyun Fan, Qing Zhang, Xiujun Lin, Yi Chen, Liu Qian, Kang Li, Xu Lu, Baodong Zheng, Lei Chen

## Abstract

This study is aim to determine the chemical composition and antioxidant activity of the essential oil from lotus leaves. The different methods and solvents were selected to extract oils from lotus leaves. About 38 components were found by GC-MS analysis, representing as 12, 15-octadecatrienoic acid (34.99%), linoleic acid and hexadecanoic acid. TBARS value, AV value and IV value reflected the various physicochemical indexes and lipid properties of *Folium nelumbinis* oil. Furthermore, antioxidant activities of the leaf samples were examined by FRAP and DPPH assays. In all systems, twice methanol-decolorized and ultrasonic-assisted essential oil using hexane solvent exhibited a higher potential activitythan than that of other extracts (ethanol, ethyl acetate and petroleum ether). These results provide a clear picture of the essential oils of *Folium nelumbinis* and demonstrate that the lotus leaves oil has an huge potential as a kind of chemical additive for the food industry owing to the strong antioxidant capacity.

## Introduction

Lotus (*Nelumbo nucifera* Gaertn) is widely known not only as a large aquatic herb but also as an edible vegetable and traditional medicine in Eastern Asia, especially in China [1]. It belongs to the Nymphaeaceae family, which is well known for its pharmacological and horticultural importance [2]. The flowers, leaves, rhizomes and seeds of lotus have been used extensively. For example, lotus seeds have a long history in using as a healthy food and also employing as a medicine to treat some ailments, including tissue inflammation, diuretic, poison antidote, emollient and skin diseases [3]. Recent years, lotus leaves are very popular in some diet tea, it has weight loss and clear heat detoxification effects, and sometimes used as food seasoning in Chinese food [4]. It also has been used in traditional medicine, stopping bleeding and resolving summer heat [5, 6].

To our knowledge, most of leaves are abandoned after the harvest of seeds and roots every year. The annual production of leaves is more than 800,000 tons, and a small amount of the leaves used as teas or traditional herbs which result in serious waste of resources and environmental pollution [7]. Therefore, the study about analysis of lotus leaves is very necessary. Catechin, myricetin-3-O-glucoside and quercetin have been reported as its major components [8-10]. In addition, few studies have examined the antioxidant of the *Folium nelumbinis* extract [11]. However, limited information is available on the *Folium nelumbinis* oil and its antioxidant activity.

The extraction process plays a significant role in the study of lotus leaves oil. Conventionally, the technologies used to extract essential oil from plants usually have some disadvantages such as low extract yield, long processing time, high temperature as traditional hydrodistillation and organic solvent extraction [12, 13] which may take some trouble. Nowadays, many new techniques have been develope to combine with organic solvents including microwave-assisted extraction, supercritical fluids, ultrasonic-assisted extraction, Ionic liquid and so on. To the best of our knowledge, no work has been published on the antioxidant activity of lotus leaves oil extracted by ultrasonic-assisted organic solvent. In the present work, the *Folium nelumbinis* from Fujian Jianning, China were used in ultrasonic extraction with different polarity solvents (n-hexane, ethanol, ethyl acetate) and compared to the soxhlet extraction with petroleum ether. Moreover, we decided to examine the antioxidant activity of each of the extract using 2-2-diphenyl-1-picryl-hydrazyl radical scavenging assay (DPPH) and ferric reducing antioxidant power (FRAP) and correlate them with Vitamin C as positive control. Thus, the composition of lotus leaves oil was identified by GC-MS and then extraction was detected, and the lipid physicochemical property was assessed including TBARS value, AV value and IV value. In addition, the antioxidant capacity of lotus leaves oil on radical scavenging activity and reducing power capacity were studied.

## Materials and methods

### Plant materials and chemicals

Leaves of nelumbo nucifera were purchased from Jianning county, Fujian province of China in summer. Ascorbic acid (vitamin C) was obtained from XiLong Scientific company. 2, 4, 6-Tris (2-pyridyl)-s-triazine (TPTZ) was purchased from Shanghai Macklin Biochemical company (Shanghai, China). 2,2-Diphenyl-1-picryl-hydrazyl (DPPH) was supplied from Sigma-Aldrich company (USA). All other reagents including n-hexane, ethanol, ethyl acetate petroleum ether (boiling range 30-60°C), methanol and DMSO were all of analytical grade, which obtained from Sinopharm Chemical Reagent Co., Ltd (Shanghai, China).

### Preparation of the plant materials

The leaves were cleaned and dried at room temperature during a night in a airy and dry environment. Then the samples (2.725 kg) were dried in 40°C for 8 h, which lost 35.78% of their water. The dried leaves were ground to the particle size of 250 μm by a mill eqiupped with three blades into powders. The weight was ultimately about 1.681kg and stocked in plastic bags in the dark until chemical analysis.

### Ultrasonic-assisted extraction

Three different solvents (n-hexane, ethanol, ethyl acetate) were used to fractionate the soluble compounds from the lotus leaves in ascending polarity [14]. Nine conical flasks (500 ml) were prepared. The samples (5 g) were mixed with 100 ml organic solvents (hexane, ethanol, ethyl acetate) in each set of three flasks. The extraction was carried out by shaking at 40°C for 3 h and subjected toultrasonic radiation using an ultrasonic water bath working at 60kHz with maximum output power of 1000 w for 25 min at 20°C. After filtration through Whatman NO.1 filter paper(7 mm), the residues were re-extracted twice again, and then the combined extracts of each sample were dried in a vacuum evaporator at 40°C. The condensed sample was further condensed with a purified nitrogen stream to 0.6 ml in volumn [15].

### Soxhlet extraction

Five grams of dry powders from lotus leaf were extracted with 100 ml petroleum ether (boiling range 30-60°C) in a soxhlet apparatus for 4 h. Then the organic solvent was evaporated by using a rotary evaporator at 40°C under reduced pressure [16].

### Once methanol-decolorized and ultrasonic-assisted extraction

The samples(5 g) were soaked in methanol(1: 20, w/v) for two groups, one was magnetic stirring 3 h at 40°C, the another was shaking 3 h at 40°C. The supernatant and sediment were separated by vacuum filtration using a Buchner funnel and 7 mm diameter size filter paper. The residues were extracted for another two times [17]. The sediments were then collected in a 500 ml baker, and 150 ml of n-hexane was added to it. After shaking in water bath for 4 h at room temperature, the sample solution was extracted by ultrasonic with powder of 100 w for 1h at 40°C. After filtration through filter paper, the solvents were evaporated by rotary evaporation at 40°C.

### Twice methanol-decolorized and Ultrasonic-assisted extraction

The sample (5 g) was soaked in methanol (1: 20, w/v) for two groups and repeated the above operation. After filtration, the sediment was collected and soaked in methanol twice again according to the same procedure. After filtration, the supernatant was then combined, evaporated and nitrogenblowed. The yellowish green powders were then stored at 4°C until application.

### Analysis of the oil

The sample diluted by DMSO (4 mg/ml) was injected into GC-MS (agilent7890GC/MSD) and then separated on an agilentHP-5ms column (length=30 m, i.d.=0.25 mm, thickness=0.25 μm). The carrier gas helium was used at a flow rate of 1.0ml/min. The ion source temperature was set at 220°C. The mass detector electron ionization was 75 eV with scanning from 20 to 600 amu. The injector temperature was set at 250°C. The gradient of oven temperaturestarted at 80°C and then raised to 300°C at 5°C/min, and was held for 20 min. The essential oil components were quantitated by relative percent peak area.

### Assessment of lipid physicochemical property

#### TBARS value

The assay of TBA was performed according to the national standard with little modification. In brief, 0.02 g of sample was mixed with trichloroacetic acid (1.0 ml) and shaking bath at 50°C for 0.5 h. After cooling to room temperature, the mixture was filtered twice with double layer filter to remove grease. And then 0.5 ml of mixture and 0.5 ml 0.3% (w/w) of thiobarbituric acid (TBA) were mixed and heated at 90°C for 40 min in a water bath. After cooling for 1 h, the absorbance was determined at 593 nm. The TBARS value was calculated from a standard curve with 1,1,3,3-tetraethoxypropane and expressed as mg MDA/kg sample.

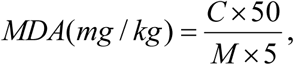

Where C is the number of micrograms of MDA found in the standard curve, µg; M is the sample quality, g.

#### Acid value (AV)

The acid value (AV) assay followed the national standard with little modification. In general, 0.02 g of sample was added to ethyl ether-isopropyl alcohol mixture (2: 1), and 2 drops of thyme phenolphthalein indicator, fully shaken and dissolved. 0.5 M standard KOH solution titrant was used to make the solution blue color. Record the volume of solution and do blank control at the same time.

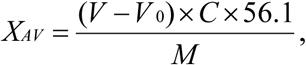

where X_AV_ is Acid value, mg/g; V is the consumed volume of the 0.5 M standard KOH solution, ml; V_0_ is the volume of the 0.5 M standard KOH solution consumed by blank determination, ml; C is the molar concentration of the standard titration solution, mol/l; 56.1 is the molar mass of KOH, g/mol; M is the sample quality, g.

#### Iodine value (IV)

The iodine value (IV) assay followed the national standard with little modification. 0.01 g of sample was added to 2 ml cyclohexane-glacial acetic acid mixture (1:1), and then added 2.5 ml Wijs reagent. After 2 h in the dark, 2 ml of potassium iodide solution and 15 ml of deionized water were added. Using 0.1 M standard sodium thiosulfate solution titrant and adding a few drops of starch solution, to titrate the solution until the iodine yellow was disappeared. Write down the volume of solution and do blank control in the meantime.

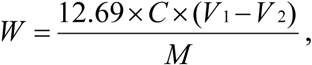

Where W is Iodine value, express as the number of grams of iodine in 100 g samples, g/100g; C is the molar concentration of the standard sodium thiosulfate solution, mol/l; V_1_ is the volume of the 0.1 M standard sodium thiosulfate solution consumed, ml; V_2_ is the volume of the 0.1 M standard sodium thiosulfate solution consumed by blank determination, ml; M is the sample quality, g.

### Ferric Reducing Ability of Plasma (FRAP) Assay

The ferric reducing ability of the extracts for each sample was measured following the previously described method with some modifications[17]. Briefly, 20 µl of sample, mixed with 180 µl FRAP reagent, freshly prepared and warmed at 37°C. The FRAP reagent contained 5 ml of 0.3 M acetate buffer, plus 0.5 ml of a 7.8 mg TPTZ powders in 40 mM HCl and 20 mM FeCl_3_.H_2_O. The samples in triplicate were added to the 96-well plate and incubated at 37°C for 20 min, then read at the absorption maximum (593 nm) by using a Microplate Reader. The Fe^2+^ solution was used to perform the calibration curve. Vitamin C at the same concentration was used as positive control.

### DPPH radical scavenging assay

The DPPH free radical scavenging activity was conducted according to a previous procedure with little modifications [18]. DPPH reagent was prepared with 0.78 mg of DPPH powders and 10 ml of pure ethanol. The mixture solution was equilibrium for 2 h. Extract solution at a rang of concentrations (0.1 ml) was mixed with DPPH reagent (0.9 ml) shaken vigorously and allowed to react 30 min at 37°C in the dark before the absorbance measuring at 517 nm. Scavenging activity of DPPH radical was calculated as a percentage according to the following equation:

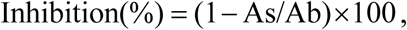

where Ab represents the absorbance without sample and As represents the absorbance with sample. Vitamin C at the same concentration was used as a reference.

### Statistical analysis

The data of all assays was expressed as standard deviation of three parallel measurements. Statistical analysis was used Duncan’s multiple range test [19] and evaluated the significant difference among various treatment with the *P*< 0.05.

## Results and Discussion

### Extract yields and Analysis

The yield of lotus leaf extracts and the essential oil extracted by four solvents (n-hexane, ethanol, ethyl acetate, and petroleum ether) are given in Table 1. The organic solvent with good solubility with samples, immiscibility with water, high extraction capability and good chromatographic behavior were considered for the extraction [20]. The yield of ultrasonic-assisted extraction by hexane, ethanol, ethyl acetate was found to be ranged from 5.74% to 6.62%, maximum extract yield by ultrasonic-assisted was obtained with n-hexane of 6.62%, and minimum with ethanol of 5.74%. Soxhlet extraction by petroleum ether was about 5.32%. According to these consequences, ultrasonic assisted hexane extraction by ultrasonic assisted took the best yield. Based on these results, the experimental results of rude oil extraction were approximately similar with other reported elsewhere. *Carthamus tinctorius L*. is a kind of Chinese old plant which extracted by n-hexane exhibiting the best yields in 2.85%, compared to the petroleum ether extraction, dichloromethane extraction and hydrodistillatin. It also exhibited the best *in vitro* inhibitory enzyme activity against PTP1B, which owned a great ability for treating obesity and diabetes [21].

**Table 1.** Extract yield percent and extract color about different extraction methods and organic solvents of *Folium nelumbinis* oil.

| Sample | Weight | Extract solvent | Extract yield % | Extract color |
| --- | --- | --- | --- | --- |
| *Folium nelumbinis* oil | 5 g powders | n-hexane | 6.62±0.55<sup>a</sup> | yellow green |
|  |  | ethanol | 5.74±0.32<sup>c</sup> | dark green |
|  |  | ethyl acetate | 6.08±0.43<sup>b</sup> | breen |
|  |  | petroleum ether | 5.32±0.41<sup>d</sup> | sepia |

<sup>a</sup>Values expressed are means ± SD. of three parallel measurements.

The extraction yield of essential oil was effected by many factors, such as solvent species, polarity, extraction time, solid-liquid ratio and so on. Among these, the solvent is the main factor that determines the extraction efficiency and composition of the extracts [22]. According to the above results, the hexane has the highest lotus leaves crude oil extract efficiency.

Ultrasound has the strong effect on the plant cells. The cells are especially tiny, fragile and with very sensitive membranes which can be broken up by sonication [23, 24]. The soxhlet extraction is a traditional method using specific techniques in the extraction of organic materials from liquid or solid matrices [25]. By using these two different techniques including soxhlet extract and ultrasonic-assisted extract with a solvent of different polarity (n-hexane, ethanol, ethyl acetate), we isolated crude extracts of sepia, yellow green, dark green and breen color. It may caused by a high total chlorophyll content (around 3000 mg/kg dissolved in cyclohexane) which led to these brown-greenish color [26].

Comparing to the extract yield and characteristic color, lotus leaves which extracted by n-hexane contained the best crude extraction. Hence, choosing the hexane extraction as the crude solvent for the next step have been determined. And concerning the comparison of these two techniques of ultrasonic-assisted and soxhlet in terms of yields and isolation times, the ultrasonic-assisted extraction was clearly fast (about 25 min), while 4 h was required for soxhlet extraction. Moreover, it is noticeable that an extraction with ultrasonic-assisted extraction for 25 min provided an appreciably high yield compared with that owned by means of soxhlet extraction for 4 h (6.62% versus 5.32%), which is the reference method about crude essential oil extraction. According to the decolorization about organic solvent methanol of different dissolution methods and docoloring times, we got the best extraction parameter of the lotus leaves essential oil in Table 2. With twice methanol-decolorized and magnetic stirring for 4 h, faint yellow transparent plant oil could be obtained to reduce the aromatic substance loss and solvent residue.

**Table 2.** Decolorizing activities about different dissolution methods and the decoloring times of *Folium nelumbinis* oil.

| Sample | Weight | Decolorization method | Solid-liquid decoloring | Number of decoloring | Dissolution method | Extract yield % | Extract color |
| --- | --- | --- | --- | --- | --- | --- | --- |
| *Folium nelumbinis* oil | 5 g powder s | Organic solvent methanol | 1:20 | once | Magnetic stirring | 1.64±0.05<sup>b</sup> | yellowish-brown |
|  |  |  |  | once | shaking | 1.68±0.04<sup>a</sup> | dark green |
|  |  |  |  | twice | Magnetic stirring | 0.89±0.01<sup>c</sup> | faint yellow |
|  |  |  |  | twice | shaking | 0.85±0.02<sup>d</sup> | dark yellow |

<sup>a</sup> Results are means ± SD. Values are the means of triplicates.

Therefore, sufficient extraction solvent of n-hexane and decolorizing solvent methanol with proper polarity could easily infiltrate to the plants and enhance the extraction and decolorization efficiency. Besides, ultrasonic-assisted extraction could be good for the penetration of solvent for enhancing the extraction efficiency and reducing extraction time.

### Chemical composition of essential oil by GC-MS

Table 3 shows the chemical composition of lotus leaves extracted by hexane. GC-MS analysis identified 38 compounds, while only 15 compounds could be primary essential oil composition. 9, 12, 15-octadecatrienoic acid (34.99%) was recognized as the main constituents, together with linoleic acid ethyl ester (12.23%), n-hexadecanoic acid (14.29%), hexadecanoic acid, ethyl ester (4.63%), and methyl (Z)-5,11,14,17-eicosatetraenoate (3.5%). In general, the most abundant components were fatty acids and esters, which were constituted mainly by octadecatrienoic acid and hexadecanoic acid.

**Table 3.**
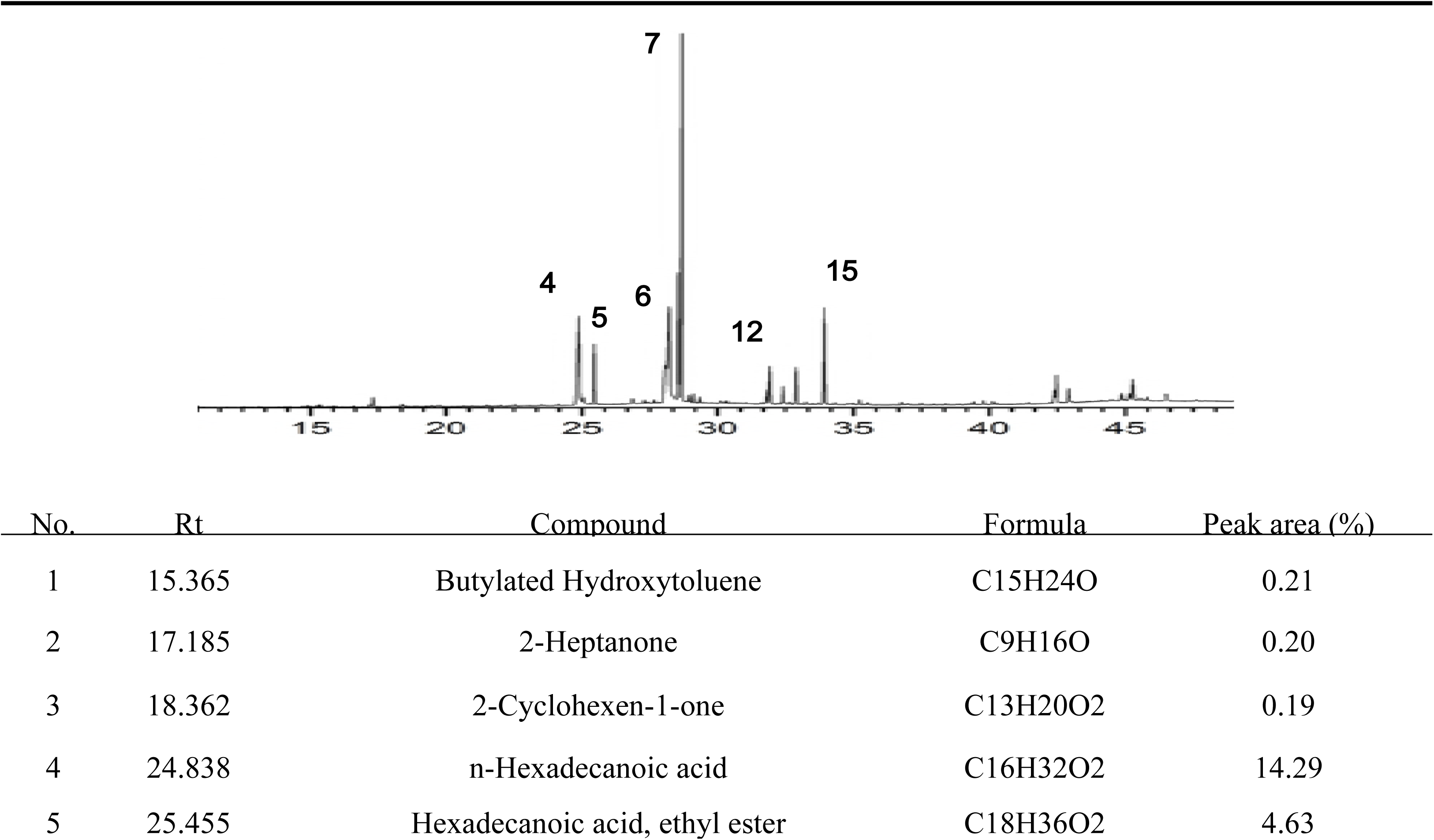
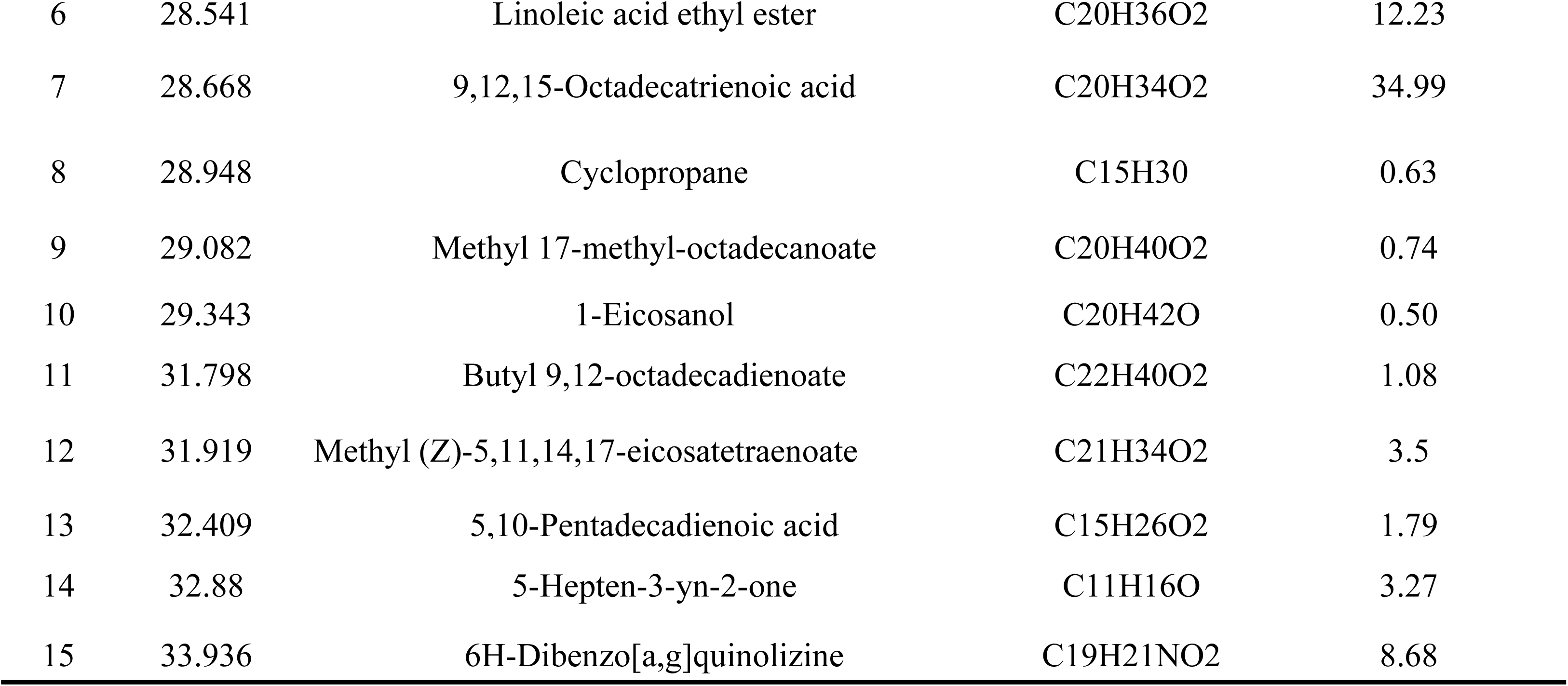
Major chemical composition of essential oil of *Folium nelumbinis*.

### Lipid physicochemical property analysis

The physical and chemical properties of oil can reflect the process quality and commodity value of plant oil, which is often applied in oil inspection. Through the current GB/T animal and vegetable oil related standards, a series of related analysis and detection were carried out on the extracted lotus leaves oil. The TBARS, AV and IV value are showed in Fig 1.

**Fig 1.**
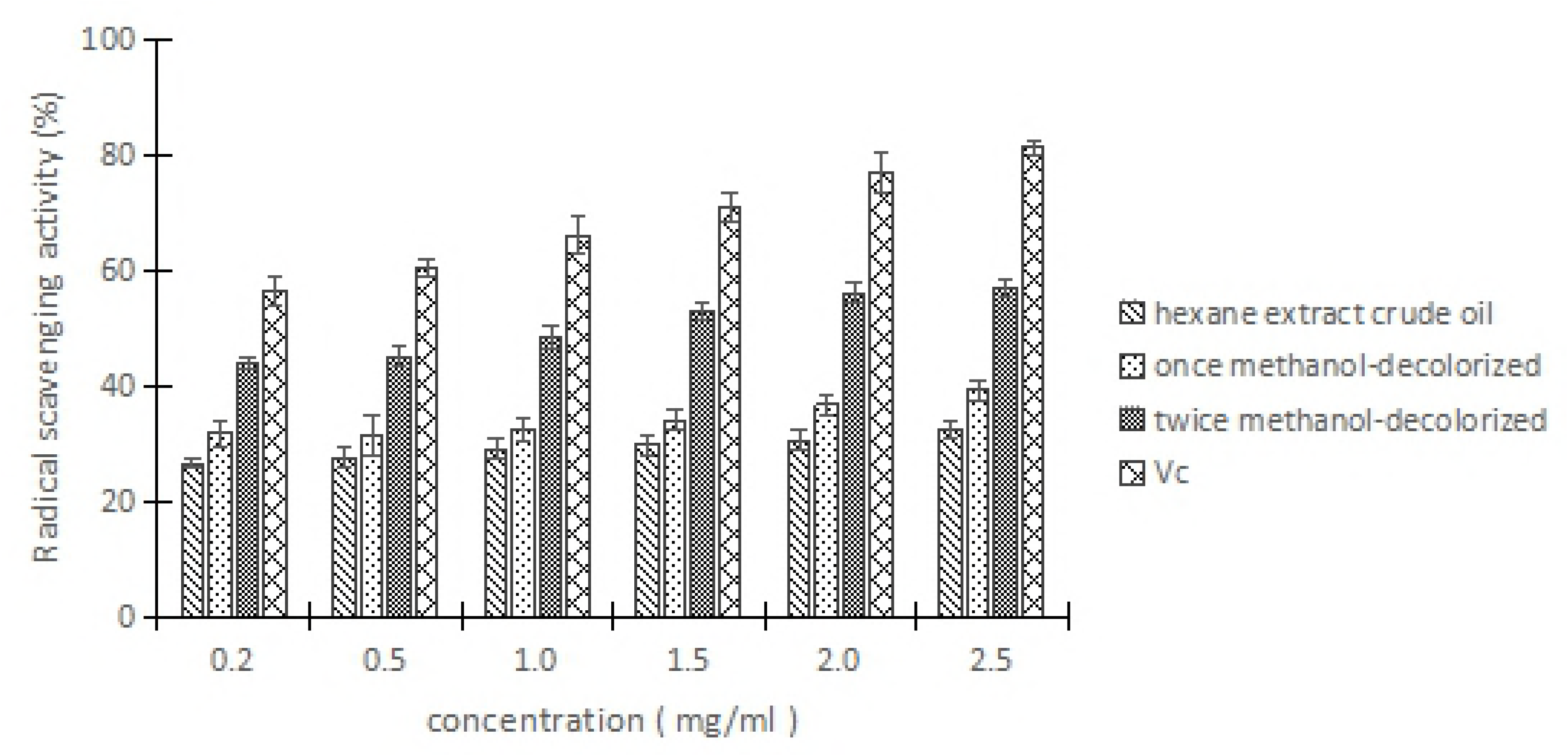
Value of TABRS, AV and IV in extraction of *Folium nelumbinis* oil. ^a^TBARS: thiobarbituric acid-reactive substances; AV: Acid value; IV: Iodine value. ^b^ Results are means±SD. Values are the means of triplicates.

Besides deeper color, lipid oxidation is a main factor to induce a decline in essential oil quality, and TBARS value is a common index of oxidative change used in oil. In the present study, storage time had an effect on lipid oxidation. On day 0, oil sample showed significantly lower value about 0.1722 of control group, oxidation level in the 150 days treated sample was significantly higher than in the control of 0 day treatment. Overall, oxidation increased with storage time of all oil samples.

Acid value (AV) is a measurement standard of the number of free carboxyl groups in fatty acids, with 1 gram of chemical substances of milligrams of potassium hydroxide (KOH). In previous studies, we found that, AV showed substantial increase of oil samples with storage time increasing. It induced the degradation of oil quality.

Iodine value (IV) is an indicator of the degree of unsaturation in oils. It is calculated by the number of grams of iodine absorbing into 100 g oils. As the same with TBARS value, AV and IV value showed significant importance with storage time. According to the IV category, lotus leaves oil may contain lots of linoleic acid and oleinic acid. It was also confirmed in the GC-MS.

### Antioxidant activities using DPPH radical scavenging

During the lipid peroxidation, free radicals play a vital role in a large amount of chronic pathologies, such as cardiovascular and cancer diseases among others [27]. DPPH is considered to be a model of stable lipophilic radical, and is one of the compounds having a free radical with a characteristic absorption at 517 nm, which decreases observably on exposure to hydrogen radical scavengers [28] and has been extensively used in estimating free radical scavenging activities of antioxidants. It is visually obvious representing as a discoloration from purple to yellow, which just like the literature said [29].

Fig 2 shows that various extractions had significant scavenging effects on the DPPH radical. Interestingly, the free radical scavenging capacity increased with the increasing extract concentration in the range of 0.2-2.5 mg/ml at a dose-dependent manner.

**Fig 2.**
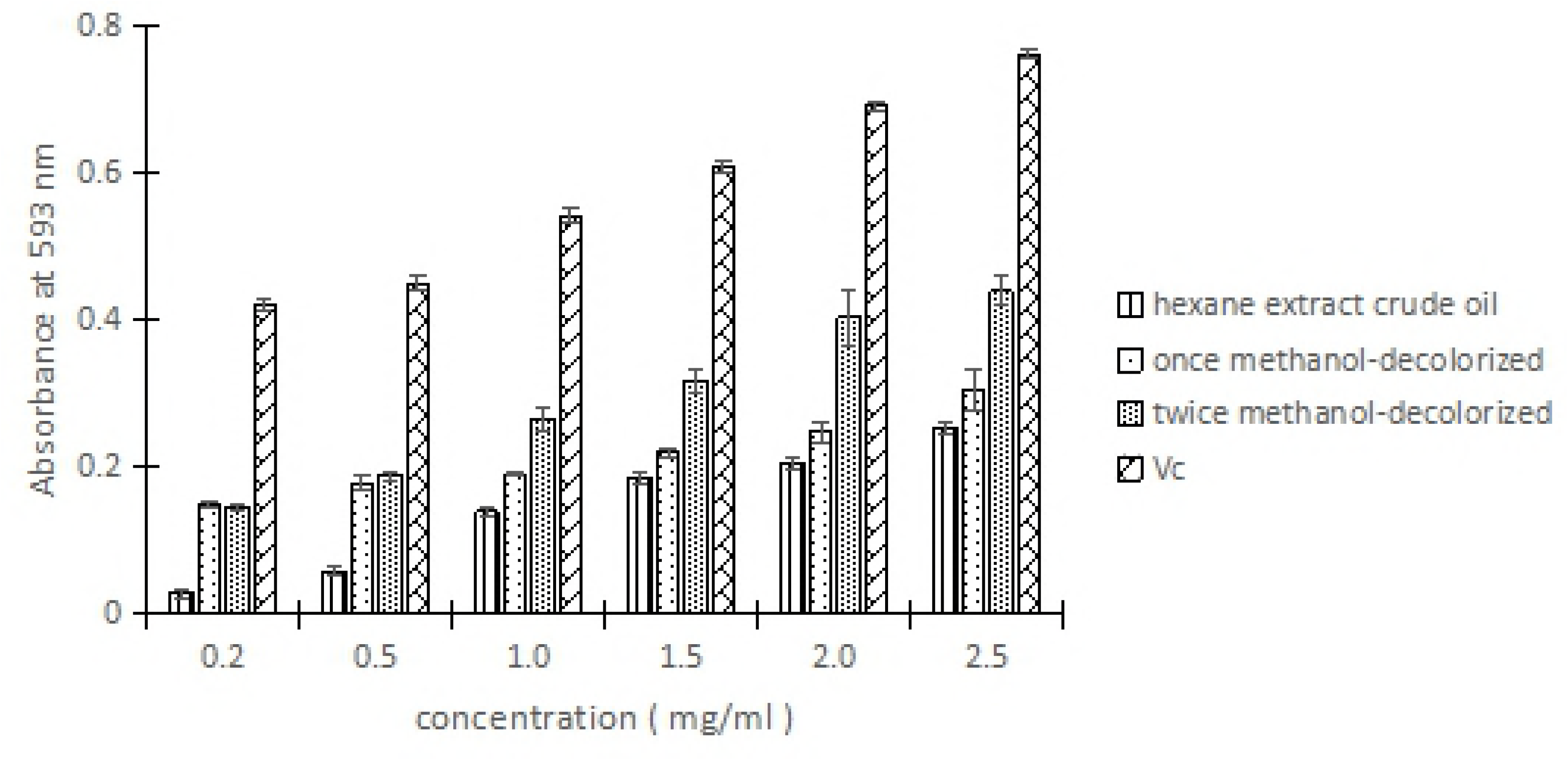
DPPH free radical scavenging activity of different methods extraction of *Folium nelumbinis* oil. ^a^Values are means of triplicate determinations (*n*=3) ± standard deviation.

Ultrasonic-assisted extracts showed a lower radical scavenging activity compared with Vitamin C, whereas the methanol-decolorized and Ultrasonic-assisted extracts showed higher percentage of DPPH. Among the two extracts by methanol decolorized, sample was only decolorized twice, which had the strongest radical scavenging activity in the extracts, and had even closer to the positive control when the concentration was at the level of 2.5 mg/ml.

These results suggested that the radical scavenging capacity of each extract was highly related to the method of extraction and the number of decolorization. The extensive investigations on antioxidant activities and free radical scavenging potential of essential oil from solvent extraction have been reported. The highest DPPH scavenging activities were showed by methanol of green tea and stem bark of *F.racemosa* when compared to other values [30]. Apart from these, the extracts of *Mentha pulegium L*. by methanol exhibited more scavenging activity than ethyl acetate and other solvent extracts [31]. The leaf of *Z. lotus* which macerated with methanol registered the high antioxidant activity [32].

### FRAP assay

In the reducing power assay, the existence of extracts induce the reduction of ferric-tripyridyl triazine complex to its ferrous form. The compound reduction capacity may be used as an important indicator for its potential antioxidant activity [33]. As showed in Fig 3, the isolated oils by different methods displayed the obvious ferric reducing power with the concentrations ranging from 0.2 to 2.5 mg/ml. Vc, as the positive control, has the powerful antioxidant effect among the test. It appears that the the high level of reducing power of Vc was always obtained, which was higher than the ultrasonic-assisted hexane extracts and methanol-decolorized extracts. The reducing power of the sample which took twice methanol-decolorized were higher than other extracts except positive control (Vc). Obviously, the reducing power of all the samples were statistically in the following order: Vitamin C > twice methanol-decolorized extracts > once methanol-decolorized extracts> crude oil extracted by ultrasonic-assisted hexane.

**Fig 3.**
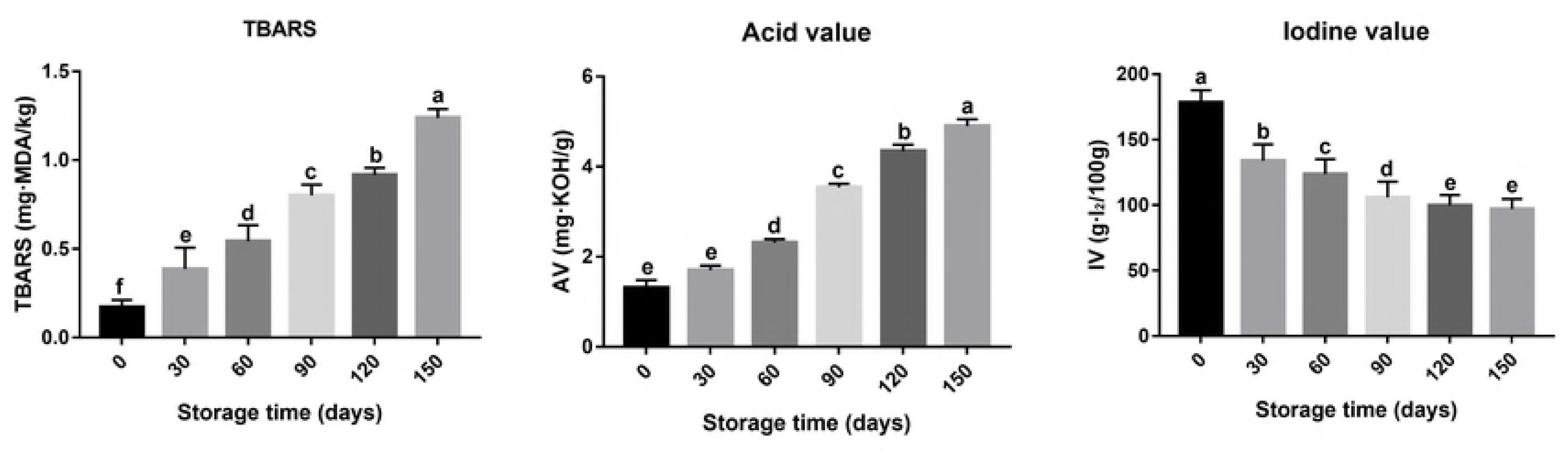
DPPH free radical scavenging activity of different methods extraction of *Folium nelumbinis* oil. ^a^Values are means of triplicate determinations (*n*=3) ± standard deviation.

It was easily found from Fig 3. that the antioxidant activity of the extract oil increased with the increase of their concentrations, and the highest FRAP value was about twice methanol-decolorized compounds with 2.5 mg/ml, thus demonstrating a great antioxidant effectiveness. This obtained result is comparable with those of natural antioxidants such as Vitamin C, namely, methanol extracts of lotus leaves are comparable to some well-known antioxidant compounds in total antioxidant system. According to these results, we can obtain that though all the extracts exhibited dose-dependent manner, methanol extracts of all the samples were found to have more antioxidant activities. The lotus leaves oil generally shows a high FRAP value indicating that it could easily donate the electron to Fe^3+^, thus reducing it to Fe^2+^[34].

## Conclusions

In summary, lotus leaves oil which extracted by ultrasonic-assisted and methanol-decolorized showed distinct characteristic in composition and antioxidant activities. Therefore, ultrasonic assisted extraction found to be the most important technology of the work. It helps the extraction of essential oil more effectively under the appropriate conditions (room temperature and pressure) in a short time. Compared to other methods, this project is simple, fast, save plant materials, lost cost and efficiency to the extraction of essential oil components. Besides, according to the antioxidant value, the *Folium nelumbinis* could be a good source of antioxidant, which suggested that the lotus leaves oil may be a high quality natural oil source and can be developed as a kind of healthcare oil. For the further studies, we need to comprehend the isolation and identification of individual essential oil compounds and also need for better understanding of their mechanism of action as antioxidant *in vivo*.

## Acknowledgments

This work is supported by the Natural Science Foundation of China (NSFC, Grant No. 31701520).

## References

1. Bo H, Ban XQ, He JS, Jing T, Tian J, et al. (2010) Comparative analysis of essential oil components and antioxidant activity of extracts of Nelumbo nucifera from various areas of China. Journal of Agricultural & Food Chemistry 58: 441–448.

2. Rai S, Wahile A, Mukherjee K, Saha BP, Mukherjee PK (2006) Antioxidant activity of Nelumbo nucifera (sacred lotus) seeds. Journal of Ethnopharmacology 104: 322.

3. Mukherjee PK, Mukherjee D, Maji AK, Rai S, Heinrich M (2009) The sacred lotus (Nelumbo nucifera) - phytochemical and therapeutic profile. Journal of Pharmacy & Pharmacology 61: 407–422.

4. Yan YL, Yu CH, Chen J, Li XX, Wang W, et al. (2011) Ultrasonic-assisted extraction optimized by response surface methodology, chemical composition and antioxidant activity of polysaccharides from Tremella mesenterica. Carbohydrate Polymers 83: 217–224.

5. Kashiwada Y,. EA, et al. (2005) anti-HIV Benzylisoquinoline Alkaloids and Flavonoids from the Leaves of Nelumbo nucifera, and Structure—Activity Correlations with Related Alkaloids. Bioorganic & Medicinal Chemistry 13: 443.

6. hui ZRGwsbydwy (2000) Pharmacopoeia of the People’s Republic of China: Chemical Industry Press.

7. Zhang L, Tu ZC, Wang H, Kou Y, Wen QH, et al. (2015) Response surface optimization and physicochemical properties of polysaccharides from Nelumbo nucifera leaves. International Journal of Biological Macromolecules 74: 103–110.

8. Ju YH, Lee K, Park J, Dong A, Shin HS (2010) Cytoprotective activity of lotus (Nelumbo nucifera Gaertner) leaf extracts on the mouse embryonic fibroblast cell. Food Science & Biotechnology 19: 1171–1176.

9. Huang CF, Chen YW, Yang CY, Lin HY, Way TD, et al. (2011) Extract of Lotus Leaf (Nelumbo nucifera) and Its Active Constituent Catechin with Insulin Secretagogue Activity. J Agric Food Chem 59: 1087–1094.

10. Lin HY, Kuo YH, Lin YL, Chiang W (2009) Antioxidative effect and active components from leaves of Lotus (Nelumbo nucifera). Journal of Agricultural & Food Chemistry 57: 6623.

11. Choe JH, Jang A, Choi JH, Choi YS, Han DJ, et al. (2010) Antioxidant activities of lotus leaves (Nelumbo nucifera) and barley leaves (Hordeum vulgare) extracts. Food Science & Biotechnology 19: 831–836.

12. Ma CH, Yang L, Zu YG, Liu TT (2012) Optimization of conditions of solvent-free microwave extraction and study on antioxidant capacity of essential oil from Schisandra chinensis (Turcz.) Baill. Food Chemistry 134: 2532–2539.

13. Suanarunsawat T, Ayutthaya WDN, Songsak T, Thirawarapan S, Poungshompoo S (2010) Antioxidant Activity and Lipid-Lowering Effect of Essential Oils Extracted from Ocimum sanctum L. Leaves in Rats Fed with a High Cholesterol Diet. Journal of Clinical Biochemistry & Nutrition 46: 52.

14. Sari?Kurkcu C, Ari?Soy K, Tepe B, Caki?R A, Abali? G, et al. (2009) Studies on the antioxidant activity of essential oil and different solvent extracts of Vitex agnus castus L. fruits from Turkey. Food & Chemical Toxicology 47: 2479–2483.

15. Yanagimoto K, Ochi H, Lee KG, Shibamoto T (2003) Antioxidative activities of volatile extracts from green tea, oolong tea, and black tea. Journal of Agricultural & Food Chemistry 51: 7396–7401.

16. Mohammed HH (2011) Soxhlet extraction of essential oil from Zingiber from officinale and optimization of process parameters.

17. Jiang SH, Wang CL, Chen ZQ, Chen MH, Wang YR, et al. (2009) Antioxidant properties of the extract and subfractions from old leaves of Toona sinensis Roem (Meliaceae). Journal of Food Biochemistry 33: 425–441.

18. Liu JR, Yang YC, Shi LS, Peng CC (2008) Antioxidant properties of royal jelly associated with larval age and time of harvest. J Agric Food Chem 56: 11447–11452.

19. Duncan DB (1955) Multiple range and multiple F tests. Biometrics 11: 1–42.

20. Sereshti H, Rohanifar A, Bakhtiari S, Samadi S (2012) Bifunctional ultrasound assisted extraction and determination of Elettaria cardamomum Maton essential oil. Journal of Chromatography A 1238: 46–53.

21. Li L, Wang Q, Yang Y, Wu G, Xin X, et al. (2012) chemical components and antidiabetic activity of essential oils obtained by hydrodistillation and three solvent extraction methods from carthamus tinctorius l. Acta Chromatographica 24: 653–665.

22. Pang J, Dong W, Li Y, Xia X, Liu Z, et al. (2017) Purification of Houttuynia cordata Thunb. Essential Oil Using Macroporous Resin Followed by Microemulsion Encapsulation to Improve Its Safety and Antiviral Activity. Molecules 22: 1697–1704.

23. Toma M, Vinatoru M, Paniwnyk L, Mason TJ (2001) Investigation of the effects of ultrasound on vegetal tissues during solvent extraction. Ultrasonics Sonochemistry 8: 137–142.

24. Vinatoru M (2001) An overview of the ultrasonically assisted extraction of bioactive principles from herbs. Ultrason Sonochem 8: 303–313.

25. Ozel MZ, Kaymaz H (2004) Superheated water extraction, steam distillation and Soxhlet extraction of essential oils of Origanum onites. Analytical & Bioanalytical Chemistry 379: 1127–1133.

26. B DE, B VV, F Rao, P Poo (2012) Characteristics of blackberry and raspberry seeds and oils. Acta Periodica Technologica 2012: 1–9.

27. Hjd D, Peltoketo A, Hiltunen R, Tikkanen MJ (2003) Characterisation of the antioxidant properties of de-odourised aqueous extracts from selected Lamiaceae herbs. Food Chemistry 83: 255–262.

28. Yamaguchi T, Takamura H, Matoba T, Terao J (1998) HPLC method for evaluation of the free radical-scavenging activity of foods by using 1,1-diphenyl-2-picrylhydrazyl. Journal of the Agricultural Chemical Society of Japan 62: 1201.

29. Sarhan, Atef M, Selim, Abdel-Hamed K, Khalel, et al. (2013) Antioxidant and antimicrobial activities of essential oil and extracts;of fennel (Foeniculum vulgare L.) and chamomile (Matricaria chamomilla;L.). Industrial Crops & Products 44: 437–445.

30. Manian R, Anusuya N, Siddhuraju P, Manian S (2008) The antioxidant activity and free radical scavenging potential of two different solvent extracts of Camellia sinensis (L.) O. Kuntz, Ficus bengalensis L. and Ficus racemosa L. Food Chemistry 107: 1000–1007.

31. Brahmi F, Abdenour A, Bruno M, Silvia P, Alessandra P, et al. (2016) Chemical composition and in vitro antimicrobial, insecticidal and antioxidant activities of the essential oils of Mentha pulegium L. and Mentha rotundifolia (L.) Huds growing in Algeria. Industrial Crops & Products 88: 96–105.

32. Elaloui M, Ennajah A, Ghazghazi H, Youssef IB, Othman NB, et al. (2016) Quantification of total phenols, flavonoides and tannins from Ziziphus jujuba (mill.) and Ziziphus lotus (l.) (Desf). Leaf extracts and their effects on antioxidant and antibacterial activities. 4: 18–18.

33. Donelian A, Carlson LHC, Lopes TJ, Machado RAF (2009) Comparison of extraction of patchouli (Pogostemon cablin) essential oil with supercritical CO 2 and by steam distillation. Journal of Supercritical Fluids 48: 15–20.

34. Meullemiestre A, Kamal I, Maache-Rezzoug Z, Chemat F, Rezzoug SA (2014) Antioxidant Activity and Total Phenolic Content of Oils Extracted from Pinus pinaster Sawdust Waste. Screening of Different Innovative Isolation Techniques. Waste & Biomass Valorization 5: 283–292.

